# Automated Maternal Behavior during Early life in Rodents (AMBER) pipeline

**DOI:** 10.1101/2023.09.15.557946

**Authors:** Hannah E. Lapp, Melissa G. Salazar, Frances A. Champagne

**Affiliations:** Department of Psychology, University of Texas at Austin, 108 E. Dean Keaton St, Austin, TX 78712

## Abstract

Mother-infant interactions during the early postnatal period are critical for infant survival and the scaffolding of infant development. Rodent models are used extensively to understand how these early social experiences influence neurobiology across the lifespan. However, methods for measuring postnatal dam-pup interactions typically involve time-consuming manual scoring, vary widely between research groups, and produce low density data that limits downstream analytical applications. To address these methodological issues, we developed the Automated Maternal Behavior during Early life in Rodents (AMBER) pipeline for quantifying home-cage maternal and mother-pup interactions using open-source machine learning tools. DeepLabCut was used to track key points on rat dams (32 points) and individual pups (9 points per pup) in postnatal day 1-10 video recordings. Pose estimation models reached key point test errors of approximately 4.1–10 mm (14.39 pixels) and 3.44-7.87 mm (11.81 pixels) depending on depth of animal in the frame averaged across all key points for dam and pups respectively. Pose estimation data and human-annotated behavior labels from 38 videos were used with Simple Behavioral Analysis (SimBA) to generate behavior classifiers for dam active nursing, passive nursing, nest attendance, licking and grooming, self-directed grooming, eating, and drinking using random forest algorithms. All classifiers had excellent performance on test frames, with F_1_ scores above .886. Performance on hold-out videos remained high for nest attendance (F_1_=.990), active nursing (F_1_ =.828), and licking and grooming (F_1_=.766) but was lower for eating, drinking, and self-directed grooming (F_1_=.534-.554). A set of 242 videos was used with AMBER and produced behavior measures in the expected range from postnatal 1-10 home-cage videos. This pipeline is a major advancement in assessing home-cage dam-pup interactions in a way that reduces experimenter burden while increasing reproducibility, reliability, and detail of data for use in developmental studies without the need for special housing systems or proprietary software.

## Introduction

Maternal behavior during early life in mammals ensures offspring survival by supporting the physical needs of the offspring including transfer of nutrients and warmth to young and protection from predators and the environment. Beyond ensuring survival, mother-offspring interactions also guide emotional, social, and cognitive development of offspring. Maternal behaviors provide stimulation to offspring through vestibular, auditory, tactile, and visual modalities during sensitive periods of elevated plasticity early in life. This somatosensory input directs development of pup physiology to have lasting effects on offspring brain and behavior across the lifespan^1–3^. Intergenerational patterns of maternal behavior are observed in several mammalian species, thus effects of these early social experiences can be perpetuated across generations^4^.

Factors in the broader ecological environment, such as chemical exposure, distal stressors, resource availability, and social context, can affect offspring development in early life through alterations to maternal offspring-directed behavior^5,6^. Maternal behavior can moderate environmental effects on offspring by buffering the effects of the environment, or increase susceptibility when disruptions to maternal care occur. Maternal effects signal to the infant the conditions of the distal environment and may modify offspring physiology to prepare the offspring for the environment they will encounter later in life^7,8^. Early environmental exposures are also potential contributors to vulnerability for neurodevelopmental disorders, which can be moderated by maternal-offspring interactions^9^.

The extensive, enduring effects of mother-offspring interactions and potential for disruption of these interactions by external influences make maternal behavior an important measure for all developmental studies. Laboratory rodent models are a preferred choice for studying the impact of early life exposures on offspring development because of the ability to precisely control environmental conditions, their relatively short lifespan, overlap in the hormonal and neural mechanisms governing maternal behavior with humans, and ability to examine the molecular physiological outcomes following early life exposures not possible or ethical in human research^10–14^. Like humans, rodent maternal behavior is influenced by the external environment and by offspring physiological and behavioral cues^15–17^. Measures of rodent pup-directed maternal behaviors typically include nest attendance, anogenital and body licking and grooming of pups, nursing (sometimes further specified into blanket, low-arched back, high-arched back, and passive nursing), nest building, and retrieval of pups when they are displaced from the nest^18^. Natural variations of home-cage maternal behaviors in Long-Evans rats are well characterized and variation in these maternal behaviors leads to shifts in offspring developmental trajectories^19–24^.

Scoring of home-cage maternal behavior live or through video recordings provides detailed information on dam-pup interactions guiding pup development^25^. However, between-study methodological variation in the acquisition and coding of this data limits comparisons between studies. Time-sampling methods, where point observations about maternal behavior are made throughout the day, permit analyses of behavior throughout the circadian cycle, but this approach has poor temporal resolution relative to focal observations and prohibits accurate quantification of behavior durations^26,27^. Frequency, duration, and bout length of maternal behaviors are increasingly used in analyses, particularly measures of “entropy” that are used to capture consistency of dam behavior patterns^28^. However, these measurements are notoriously time-intensive to score from videos and involve substantial training on coding maternal behavior. To address these issues, more sophisticated tools are needed to quantify maternal behavior over longer periods of time.

Recent advances in machine learning tools for behavior analysis have the capacity to build standardized behavior pipelines for video data to produce high-quality and highly-reproducible measurements^29^. Using open-source tools, we developed the Automated Maternal Behavior during Early life in Rodents (AMBER) pipeline for scoring home-cage maternal behavior. AMBER uses DeepLabCut^30,31^ to extract pose estimation data from rat dam and pups from home-cage recordings within standard cages. Side-view recordings permit tracking of key body points on dams and pups, unlike top-down recordings where the dam occludes pups during nest attendance. AMBER then uses Simple Behavior Analysis^32^ (SimBA) to run behavior classifiers for seven maternal behaviors: nest attendance, licking and grooming, active nursing, passive nursing, self-directed grooming, dam eating, and dam drinking. We applied the AMBER pipeline to a set of 242 videos from postnatal day (P) 1-10 to evaluate maternal phenotype. All scripts and models used in the pipeline are publicly available (https://github.com/lapphe/AMBER-pipeline; https://osf.io/e3dyc/).

## Results

### Video recording

Home cage behavior was recorded from Long-Evans dams and litters on postnatal day (P) 1-14 with Raspberry Pi 3B+ minicomputers equipped with Raspberry Pi Module 1 NoIR cameras. One hour and 24-hour recordings captured video during the light and dark phase under infrared LED lights.

### Dam pose estimation model

Pose estimation for dams was achieved using single animal DeepLabCut^30^. 4,710 frames were extracted from 255 videos. Thirty-two dam body points were labeled providing adequate coverage of the dam regardless of partial body occlusion or body orientation relative to the camera (**Figure 1B**). Two percent of labeled frames were used for a test set for model evaluation and the remaining 98% of labeled frames were included in the training set. The dam model was trained for 650,000 iterations saving a snapshot every 50,000 iterations using ResNet-50 with a batch size of 8. Model performance was evaluated at all snapshots and loss, a quantification of the error between predictions and true values (user labels) during training, was calculated every 1000 iterations (**Figure 2A**). Snapshot 10 (550,000 iterations) had the best performance (comparing model-predicted key point location to user key point annotation location) on test frames on test frames with an average error of 14.39 pixels (4.1–10 mm depending on animal depth in frame) and 6.36 pixels on training frames after filtering to only points with a likelihood cutoff threshold above 0.5 (**Figure 2B**). Visual inspection of labeled held-out videos confirmed model performance.

**Figure 1.**
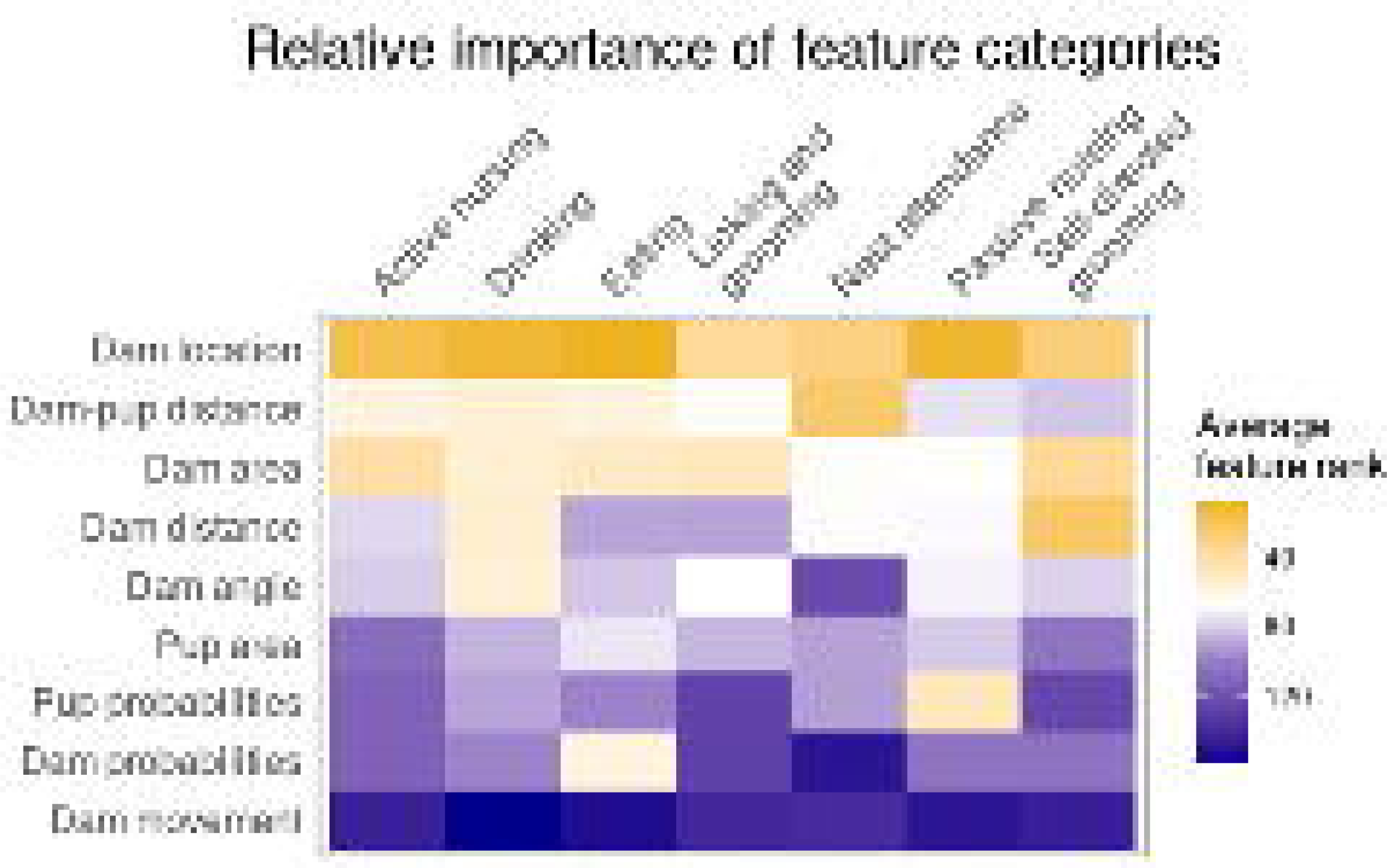
A) AMBER pipeline overview. Videos can be recorded with any device that provides sufficient resolution to visualize the dam and pups clearly. Videos can be optionally preprocessed to convert to greyscale, reduce resolution, or cropped to reduce downstream computational processing time. After video recording, pose estimation is accomplished using trained networks to detect dam and pup key points with single animal DeepLabCut and multi-animal DeepLabCut respectively. Next, coordinate pose estimation data for dams and pups is joined in one csv file. SimBA behavioral classifiers trained to detect seven maternal behaviors are run on the pose estimation data to produce frame-level behavior annotations. B) Recording set up. Raspberry pi cameras are pictured set up at the nest end of the cage to capture side-view recordings. C) Dam and pup pose estimation key points. The dam pose estimation model is trained to detect 32 key points on dams. The pup multi-animal model is trained to identify nine key points on each individual pup visible.

**Figure 2.**
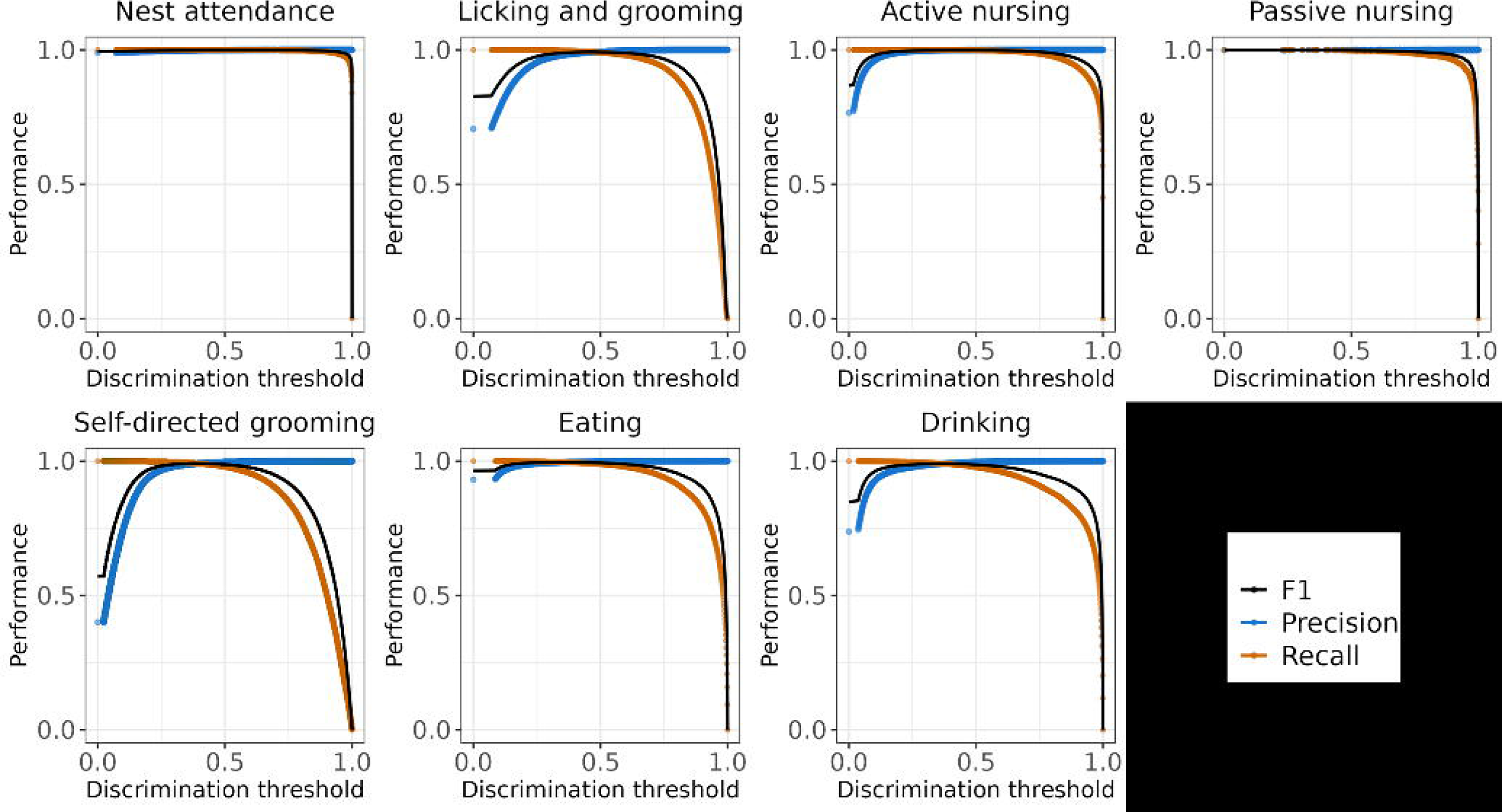
A) Dam pose estimation and C) Pup pose estimation model training statistics. Loss, a measure of model performance during training, decreased with each training iteration. B) Dam and D) Pup pose estimation model snapshot evaluations. Average root mean square error (RMSE) in pixels across all key points is plotted for training and test sets with and without a probability cutoff of 0.5 at each snapshot during model training.

### Pup pose estimation model

The pup pose estimation model was developed using multi-animal DeepLabCut^31^. 1,712 frames from 238 videos were extracted using k-means clustering and uniform distribution. Nine pup points from the nose to tail base were labeled on each pup visible (**Figure 1C**). Two percent of labeled frames were reserved for model evaluation and the remaining 98% of labeled frames were included in the training set. The pup model was trained for 200,000 iterations saving a model snapshot every 5,000 iterations using DLCRNet_ms5 with batch size of 8. Model loss was calculated every 1000 iterations (**Figure 2C**). Pup pose estimation model performance was evaluated at all snapshots (**Figure 2D**). The final snapshot had the best performance, with an average error of 4.67 pixels on training frames and 11.81 pixels (3.44-7.87 mm depending on proximity of pup to camera) on test frames after filtering to points with a likelihood cutoff threshold above 0.5. Root mean square error (RMSE) for individual pups and each body point was similar (**Supplemental Figure 1**). Visual inspection of labeled hold-out videos confirmed accurate tracking of individual pup body parts for pup detections. While pups remained in the nest the majority of the time, pups outside the nest were also tracked provided a sufficient portion of their body was visible in the frame.

### Pose estimation data postprocessing

Pups are often in a huddle mass in the nest in the home-cage, with many pups fully or partially occluded by other pups, the dam, or nesting material at any given time. This occurrence presents a challenge for assignment of pup body part detections to individual pup identities during individual assembly (tracklet creation and tracklet stitching) across frames in the second half of the multi-animal DeepLabCut workflow. Although pup detections achieved good performance, we observed substantial loss of pup points after individual assembly, where only pups with a majority of tracked body points visible had reliable tracking. A primary goal of tracking pups for this pipeline is to identify nest location, so the litter can be treated as a unit rather than identify individual pups. Therefore, all pup key point detections, at the midpoint of the multi-animal workflow by running the DeepLabCut function analyze_videos with auto_track= false, rather than individual pup tracks obtained at the end of the workflow, were used for pup pose estimation. This ensured that all pup detected key point coordinates are kept for the entire litter, with the drawback of not know which points belong to specific pups. A custom python script (pheno_pickle_raw.py) was used to convert the pup detections pickle file to a csv file containing pup key point detections. Raw detection key points assigned to individual pups after conversion to csv do not necessarily belong to the assigned pup.

Next, unfiltered dam and pup pose estimation files were joined by frame number using a custom script (join_dam_pup.py). Column headers were reformatted to match the expected input by SimBA.

### Behavior classifier project set up

Simple Behavior Analysis (SimBA) was used to generate seven behavior classifiers from pose estimation data from a subset of videos using the standard workflow except where noted below (https://github.com/sgoldenlab/simba)^32^. We used a single animal SimBA project configuration with user-defined body points consisting of all dam and pup key points. Width of the wire cage top at the lowest point (approximately half the depth of the cage) was used to define pixels per mm during the Video Settings step (**Supplemental Figure 3**). The outlier correction step of SimBA was skipped because it relies on body-length distance across frames to perform these calculations, which is influenced by the dramatic differences in body length when the dam is near the front versus back of the cage. Instead, low probability key points are accounted for during feature extraction by weighting calculations by key point probabilities.

### Classifier feature extraction

SimBA behavior classifiers train on features derived from pose estimation data. Dam and pup features were extracted from the pose estimation data using a custom script to create dam-specific and pup-specific features (**Supplemental Table 1**). Features were derived from pup pose estimation data (19 features), dam pose estimation data (172 features) or both dam and pup data (27 features). Features can also be broken down into categories: dam location (e.g. y coordinate of dam centroid), dam areas, dam key point angles, dam key point probabilities, dam movement, pup area, pup probabilities, and dam-pup distances. Features also included summary statics (mean, sum, standard deviation) across rolling windows of .1s, 1s, and 2s. In addition, 30 m and 60 m rolling windows for pup centroid were calculated and used for some dam-pup distance features to account for longer periods when pups are mostly or completely occluded (e.g. by bedding or the dam).

### Random forest behavior classifiers

Classifier training videos were carefully annotated for seven maternal behaviors using BORIS^33^ then imported into SimBA. A total of 3,366,254 frames (31.1 hours of recording at 30 fps) from 28 one-hour videos and 10 additional shorter clips (1.5 −22 m each) were annotated for nest attendance, active nursing, passive nursing, licking and grooming, self-directed grooming, eating, and drinking. Behavior definitions are provided in **Figure 3** and a detailed ethogram of these behaviors can be found at https://github.com/lapphe/AMBER-pipeline^16,17^. Because of the large number of frame annotations, the frames used for training classifiers was reduced by taking every other frame from each video. Adjacent frames are likely to have similar features and behavior annotations, so this allowed for a reduction in data set size while maintaining diversity of the training set within and across videos. 20% of remaining frames were used as a test set to evaluate model performance. Frequency of behaviors in the training and test set ranged from 2.5% (passive nursing) to 47.8% of frames (nest attendance). Random forest models were run in SimBA with the following hyperparameters: 100-1500 trees, minimum node = 1 or 2, RF_criterion = gini, RF_max_features= sqrt, test size= 20%, and no sampling adjustment (**Supplemental Table 2**).

**Figure 3.**
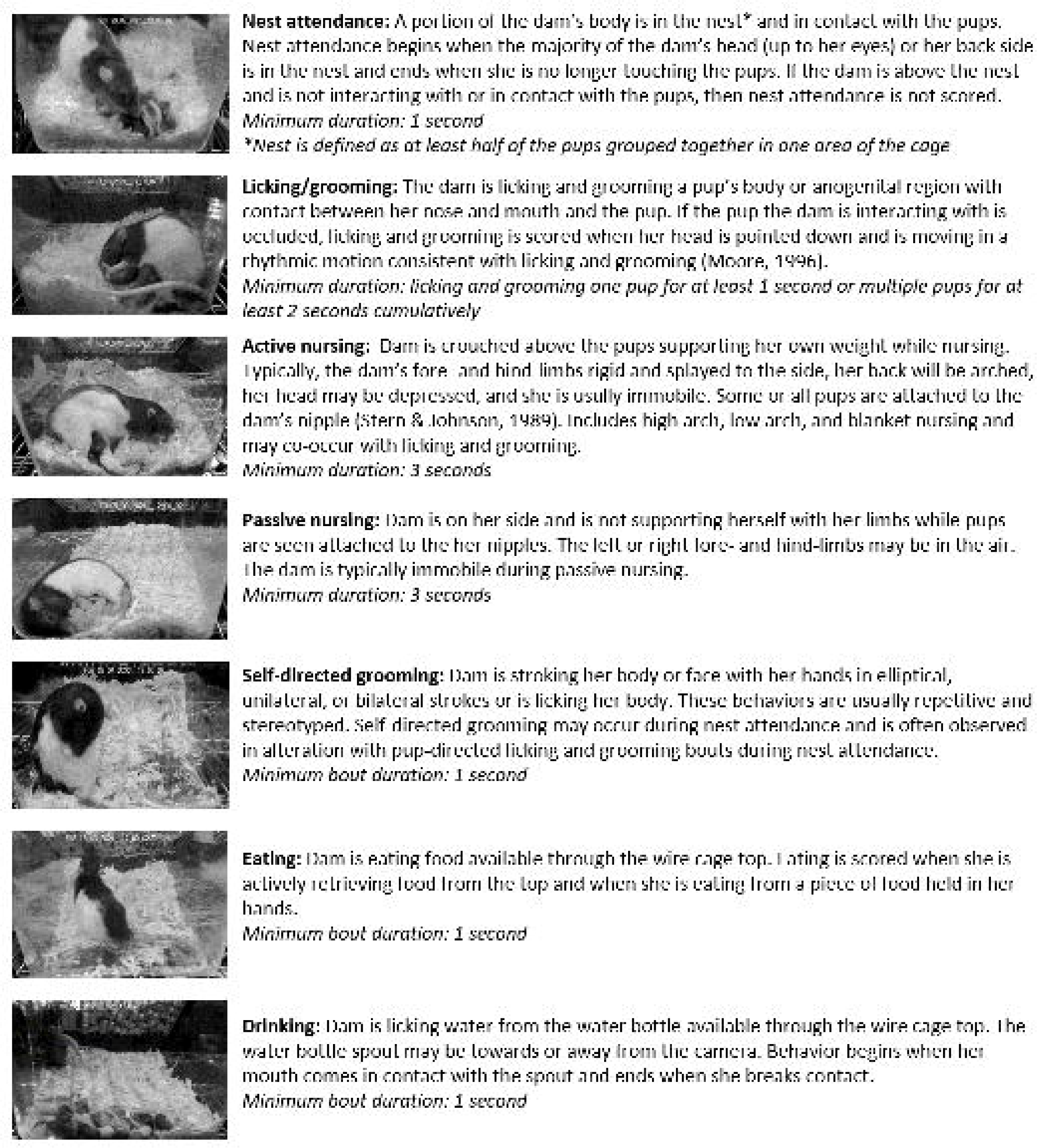
Brief behavior descriptions for each of the seven maternal behaviors annotated to train the SimBA behavioral classifiers. Additional ethogram information can be accessed at https://github.com/lapphe/AMBER-pipeline.

An additional four one-hour recordings (30 fps) were manually scored as a hold-out video validation set. Because no frames from the hold-out videos are used for model training, this allows for evaluation of model generalizability.

### Behavioral classifier evaluation

Behavior classifiers were evaluated by calculating the precision (fraction of true positives among all frames scored as positive), recall (fraction of true positives retrieved out of all true positives in data set), and F1 scores (harmonic mean of precision and recall) for all models (**Supplemental Table 2**). All behavior classifiers obtained good accuracy at or above 0.886 on the test fraction of frames. Discrimination thresholds, or probability above which a behavior is classified as present, were determined using precision-recall curves and visually inspecting behavior predictions in videos (**Figure 4; Supplemental Table 2**). Classifier performance on the hold-out video set remained high for nest attendance (F_1_ =.990), active nursing (F_1_ =.828), and licking and grooming (F_1_ =.766). Self-directed grooming, eating, and drinking classifiers performance was substantially lower (F_1_ =.534-.550). Lower performance for eating and drinking was partly due to overlap in false positive and false negatives between these classifiers. Considering eating and drinking as a single behavior improved performance (eating or drinking precision = .71, recall =.69, F_1_= .70). Passive nursing did not occur in the hold-out video set, precluding performance evaluation for that classifier.

**Figure 4.**
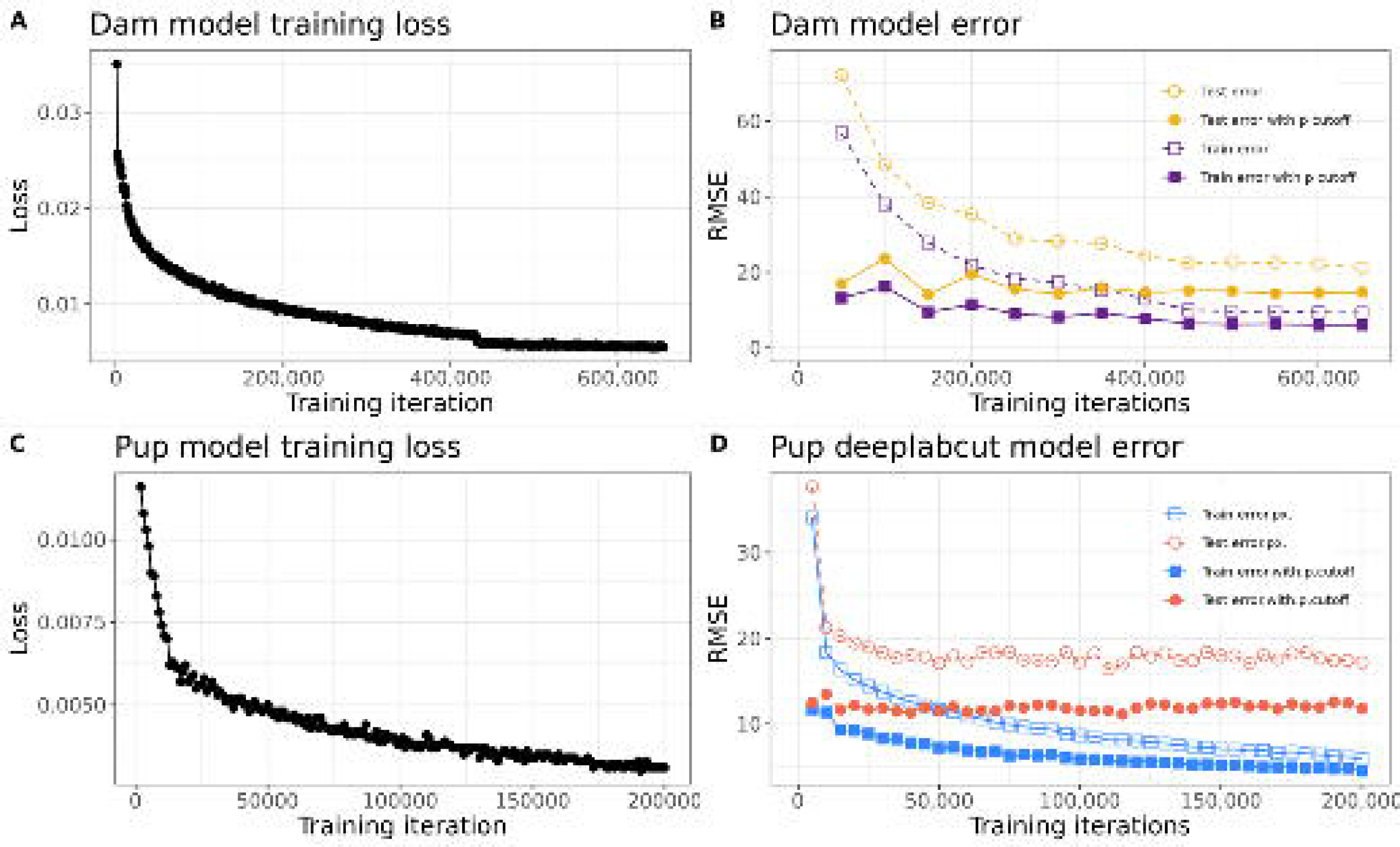
Accuracy-discrimination threshold curves for SimBA classifiers. Discrimination thresholds are cutoff values for determining if a behavior is present. Precision (accounts for false positives), recall (accounts for false negatives) and F1 scores (harmonic mean of precision and recall) at different discrimination are shown for all behavior classifiers.

### Pipeline validation for maternal phenotype

A set of 242 videos of one-hour home cage recordings taken on P1-10 from 49 dams were analyzed with AMBER pipeline workflow as shown in **Figure 1A**. Thresholds used for behavior classifiers are noted in **Supplemental Table 2**. Total duration, percent time, bout number, and mean bout duration were calculated in SimBA for each behavior for each video (**Figure 5** and **Supplemental Figure 5**). Change in behavior durations over time was analyzed using linear mixed models in R with the lmerTest package with litter ID included as a random effect^36^. Nest attendance (β = −179.61, t = −10.12, *p*< .001), licking and grooming (β = −18.37, t = −4.11, *p*= .01), active nursing (β = −120.29, t = −7.93, *p*< .001), and passive nursing (β = −2.15, t = −2.17, *p*=.03) significantly decreased with litter age. Drinking (β = 33.64, t = 7.15, *p*<.001) and eating (β = 18.68, t = 3.05, *p*<.01) significantly increased with litter age and self-directed grooming (β = 9.649, t = 1.42, *p*=.15) did not significantly change with litter age (**Figure 5**). The mean percent time engaging in the behavior across all videos was 39.3% for nest attendance (min = .3%, max = 100%, median = 32.7%), 10.1% for licking and grooming (min=0, max = 32.2%, median = 9.4%), 21.6% for active nursing (min = 0, max = 98.4%, median = 16.59%), 0.2% for passive nursing (min = 0, max = 11.1%, median = <1%), 16.85% for self-directed grooming (min = 0, max = 66.9%, median = 15.6%), 9.5% for eating (min = 0, max = 59.7%, median = 7.1%), and 7.0% for drinking (min = 0, max = 43.2%, median = 5.4%).

**Figure 5.**
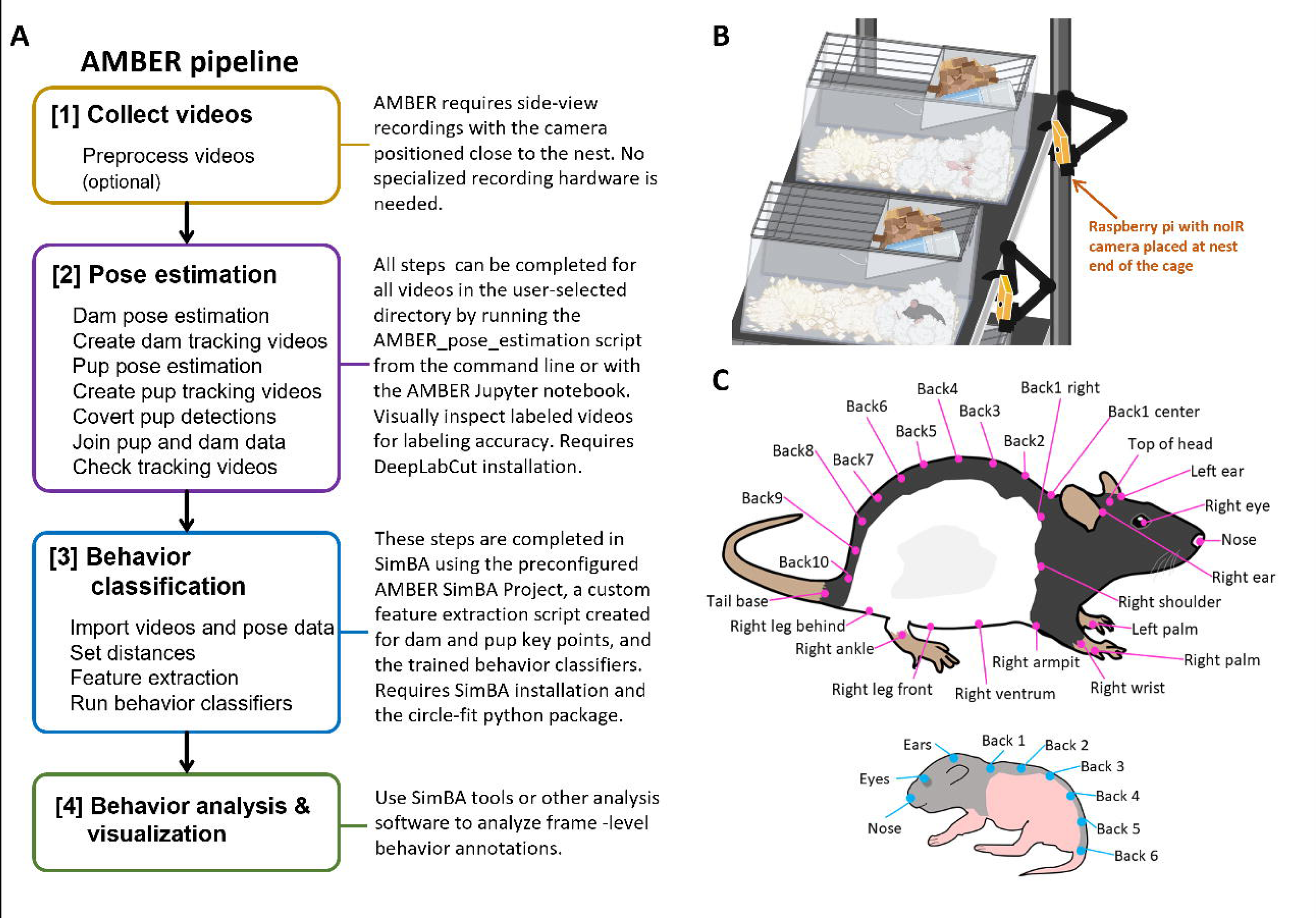
AMBER-scored maternal behavior measures. A) Total behavior duration over postnatal days. Total duration of nest attendance, licking and grooming, and active nursing during the one-hour recordings decreased as litter age increased. Eating and drinking increased as litter age increased. B) Distribution of percent time engaged in each behavior for each video. Percent time dams spent on the nest, licking and grooming, and engaging in self-directed grooming was relatively normally distributed.

### AMBER pipeline deployment

Although AMBER relies on the capabilities of DeepLabCut and SimBA software, it deviates significantly from the standard workflows and involves additional steps to work. To improve user experience and reduce barrier of entry for inexperienced programmers, we provide materials to simplify the workflow and reduce user burden for large video sets (**Figure 1A**). First, given that DeepLabCut is installed, all pose estimation steps can be performed automatically with a single command line function using the AMBER_pose_estimation.py runner script (e.g. python AMBER_pose_estimation.py path/to/videos). This program performs dam and pup pose estimation, converts pup detections, combines dam and pup data, and formats data for all videos in the indicated video directory. Alternatively, users can perform the same steps or modify code using the provided Jupyter notebook. Next, the pre-configured AMBER SimBA project can be used to perform the behavior classification steps. Instructions for implementing the AMBER pipeline are available at: https://github.com/lapphe/AMBER-pipeline.

### Post-hoc explanability metrics for behavior classifiers

While not part of the AMBER pipeline, explanability metrics offer interpretable descriptions for how model decisions are made from feature values^34^. Feature importance permutations provide an estimation of information loss when the feature is replaced with randomly shuffled values from the same distribution as the original feature data. Feature importance permutations were calculated for each behavior classifier with eli5 python library in SimBA. Relative importance of each feature within each model was determined by ranking features from most important (rank 1) to least important (rank 217) based on feature importance score for between-model comparison. The average rank for each feature category was calculated for heatmap visualization (**Figure 6**). Dam location features are the most important feature category (lowest average rank) for active nursing, drinking, licking and grooming, passive nursing, dam-pup distance features were most important for nest attendance, and dam key point distances were most important for self-directed grooming. Dam movement is the least important average feature category (highest rank) for all models except the nest attendance.

**Figure 6.**
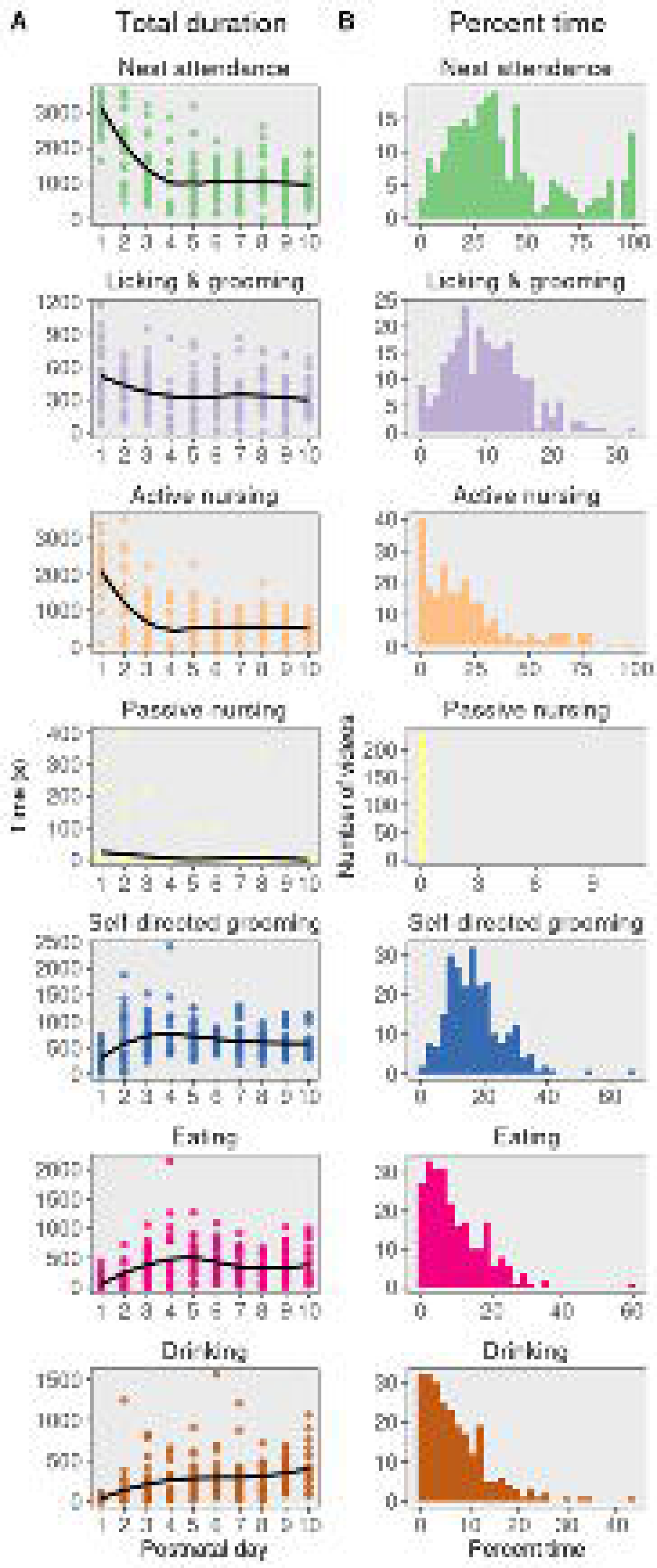
Relative importance of feature categories as determined by feature importance permutations. All features were ranked from most important (rank 1) to least important (rank 218) based on mean feature importance score for each behavior classifier. The title color corresponds to the average rank of all features belonging to each category. Yellow tiles are indicative of more important feature categories (rank closer to 1) and dark blue tiles are indicative of less important feature categories (rank closer to 218).

Shapely Additive Explanations (SHAP) is another explanability metric that uses a game theoretic approach that can be applied to tree-based machine learning models to allocate the contributions of individual features to the overall final behavior probability in a frame based on magnitude of feature attributions^34,35^. SHAP analysis was run in SimBA on 150 random frames with behavior present and 150 random frames with behavior absent for each model to calculate individual feature contributions to overall frame behavior probability (https://github.com/slundberg/shap). **Figure 7A** shows the top six features with the largest absolute SHAP scores for each behavior classifier, where the solid black line is the base rate for the behavior (probability of a given frame containing the behavior by chance), each individual point reflects the change in behavior probability relative to the base rate (SHAP score) for that feature for one frame, and the color of the point reflects the z-score of the actual feature value for that frame. Consequently, the relationship between behavior probability shift and actual feature value can be deduced as positively or negatively associated. For the nest attendance classifier, features with high SHAP scores include distances between dam centroid and pup centroid and dam convex hull features. Likewise, classifiers for other on nest behaviors (active nursing, licking and grooming, passive nursing) also include dam-pup distance features and dam convex hull features among the top SHAP features. Top features for self-directed grooming and licking and grooming classifiers include ear movement and dam distances. The sum of SHAP score for all features by feature categories (**Figure 7B**) show that dam movement features did not have a substantial impact on nest attendance, but had a moderate influence on increasing behavior probabilities in remaining models and had a particularly large effect on licking and grooming, self-directed grooming, and active nursing. Pup probabilities and pup area features had little effect on behavior probabilities.

**Figure 7.**
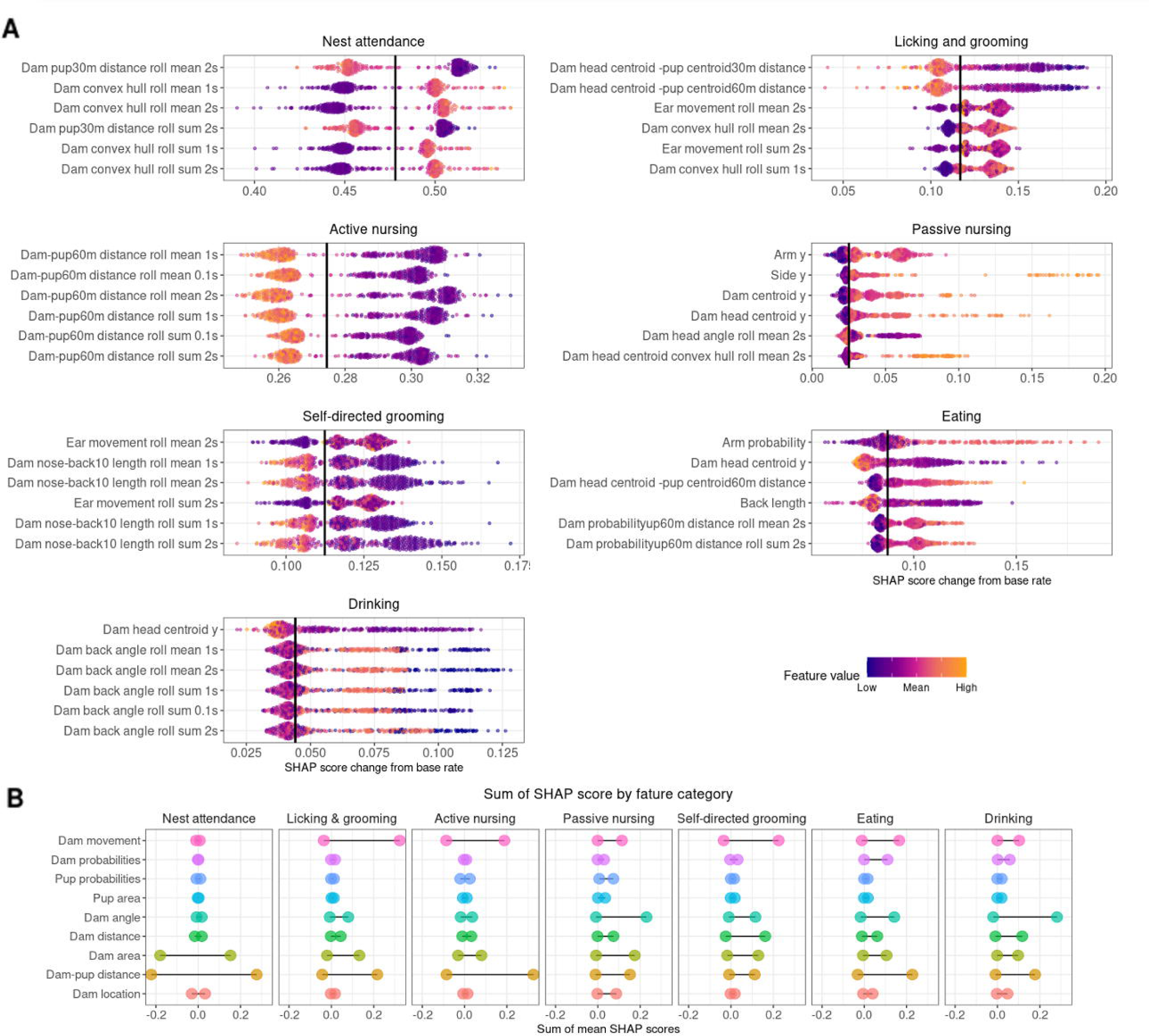
SHAP results. A) Top six features with highest SHAP scores for all models. SHAP scores with the largest average absolute value across all tested frames are shown for each behavior classifier. Each point denotes the SHAP score for one of the 150 frames included in SHAP analysis. The solid black line shows the base rate for each classifier, or the probability that the behavior occurs given the frequency of behavior in the training data set before consideration of feature information. Base rates vary by classifier and are shown in Supplemental Table 2. SHAP values greater than the base rate indicates positive SHAP scores for that frame, increasing the behavior probability. SHAP scores to the left of the black line indicate scores that reduce the behavior probability in that frame. Each point is colored by the z-scored value of the feature for that specific frame, indicating the relationship between feature values and shift in behavior probabilities. B) Sum of SHAP scores of all features in each feature category for all classifiers. Positive SHAP scores indicate the magnitude of the feature influence on increasing the behavior probability when the behavior is present and negative SHAP values indicate influence on reducing the behavior probability in frames where the behavior is absent. 0 represents the base rate of behavior in the training dataset.

## Discussion

Continuous home-cage monitoring is an optimal approach to assess dam-pup interactions in a laboratory setting, but the burden of manual scoring limits the implementation of this approach. We present a pipeline that automates scoring of rodent dam-pup home-cage video recordings to produce frame-level annotations of seven maternal behaviors with high accuracy. Pup-directed maternal behaviors performed particularly well on the hold-out video set with F_1_ scores of .990 (nest attendance), .828 (active nursing), and .766 (licking and grooming). AMBER uses open-source software, standard rat housing equipment, and does not require any specialized recording hardware or animal identification markers. When paired with automated recording equipment, home-cage behavior can be collected from an entire cohort of animals simultaneously for long time periods while avoiding the effects of experimenter presence or bias on behavior. Maternal behavior affects a variety of developmental outcomes, and AMBER eliminates the reproducibility concerns, training, and inter-rater reliability drawbacks of manual scoring home cage maternal behavior, allowing assessment of maternal behavior regardless of behavior expertise.

The validation set of recordings show that AMBER-scored videos produce expected patterns of maternal behaviors over the first ten postnatal days. The duration of pup-directed behaviors was high on P1 and declined over the first week while the durations of eating and drinking increased. This is consistent with previous work from manually-coded behavior reporting declines in dam-pup contact and licking and grooming from P1-10^37^. Licking and grooming, nest attendance, and self-directed grooming were relatively normally distributed, with the range of percent of time for licking and grooming similar to frequency observed in time-sampling studies for Long-Evans rats^37^. Active nursing was not normally distributed, although this difference may be attributed to the one-hour observation during early dark phase for the present study versus sample observations throughout the dark and light cycle used in previous work^38^. The number of bouts for behaviors was very high in a few videos (**Supplementary Figure 3**) and mean bout duration was lower than expected based on manually scored data from previous studies. This difference is explained by frame-level behavior measurements, where one frame labeled with the behavior presence (or absence) is sufficient to determine a “bout”. Smoothing methods such as employing a minimum bout duration can be applied directly in SimBA to filter the data.

Using the AMBER pipeline as presented in **Figure 1A**, users get frame-level resolution of seven maternal behaviors. However, the individual components of AMBER (dam pose estimation, pup pose estimation, and behavior classification) can also be used for other applications. First, dam pose estimation and pup pose estimation models may also be used separately in other contexts to track adult rats and pups in any side-view recording which is more compatible with most standard rodent home cages. Second, pose estimation data may be used with compatible software to perform unsupervised behavior clustering^39–41^. Third, specific features, e.g. the convex hull area of pups, dam centroid-pup centroid distance, or degree of dam back curvature during nursing, extracted from the pose estimation data may be informative in concert with behavior annotations. These data are calculated during feature extraction and are readily available for further analysis. Finally, the small size of neonatal pups and large number of pups in the litter makes manual behavior scoring for pups very difficult. While the primary purpose of the pup pose estimation model in the AMBER pipeline is to determine pup and nest location and dam-pup distances, pup pose estimation data could be used separately to evaluate pup behavior in relation to dam behavior.

Model explanability metrics shed light on the “black-box” of behavioral classification by providing articulated descriptions of how features influence model performance that allows users to critique model construct validity and compare different models beyond model performance^34^. Feature importance permutation results showed low importance of dam movement features. The majority of features comprising the dam movement category are the movement of individual dam body points, so this may suggest that this information is less informative or reliable for behavior classifier predictions than features that use information from multiple body points. SHAP analysis revealed several intuitive relationships between dam pup-directed behaviors and change in behavior probability in our classifiers: 1) the convex hull areas of the dam will get larger as she moves toward the nest (closer to the camera) at the front of the cage; 2) dam-pup Euclidian distances will decrease when the dam is interacting with pups and will increase when the dam is off-nest; 3) dam movement features are informative for classifiers that can be operationally defined by specific body movements (e.g. licking and grooming); 4) the water bottle is located in the top of the cage, so the angle of the dam’s back can be informative in identifying drinking behavior.

AMBER is a substantial improvement over manual scoring methods for dam-pup home cage behavior, but also has some notable limitations. Classifier performance for pup-directed behaviors may be compromised in recordings where pups are occluded for the duration of the video since pup coordinate information and dam-pup distances are important features for several classifiers. This shortcoming could be circumvented by manually adding nest location information in place of pup tracking. Furthermore, the behavior classifiers presented here are trained on tracking information from side-view recordings and will not generalize to top view recordings. We chose side view recordings to allow for better pup tracking and to eliminate the need for any specialized home-cage equipment as many standard home cages contain food and water in the cage lid. In addition, F1 scores for self-directed grooming, eating, and drinking classifiers were lower on the hold-out video set compared to the test set. The improvement in performance when combining eating and drinking behaviors suggest that information about the location of the food and water could improve the models. Finally, the pose estimation models at present are optimized for detecting key points in Long-Evans rats and are unlikely to generalize well to other rodent species that are visually different without training on additional frames. Likewise, differences in camera angle, bedding material, enrichment objects, cage layouts, or lighting in recordings compared to the training videos may interfere model transferability, requiring some additional labeled frames and pose estimation model retraining^42^. We are currently expanding these model training sets to include frames from videos of Sprague-Dawley rats, C57/Bl6 mice, and CD1 mice in different home cages to make the pose estimation models more robust and able to perform well for a wider variety of rodent developmental studies. These models will be made publicly available on the AMBER repositories. Despite these current limitations, the AMBER pipeline is a significant step forward for improving analysis of home cage dam-pup interactions to provide standardized, detailed behavioral data likely to yield new insights in developmental studies.

## Materials and methods

### Animal husbandry and breeding

All animal protocols were approved by the IACUC at the University of Texas at Austin, were performed in accordance with IACUC guidelines and regulations, and are reported in accordance with the ARRIVE guidelines. Animals were housed in polycarbonate cages (19” × 10.5” × 8”) with standard wire tops and were kept on a 12:12 h light cycle (lights off at 10 am EST). All dams were provided with Aspen shavings (Nepco) for bedding material, which can be manipulated by dams to construct nests. No additional bedding was provided. All animals were fed standard chow (Lab diet 5LL2) and water *ab libitum* through bottles held at approximately a 45-degree angle in the wire tops. Eighty-eight adult P60–70 Long-Evans females and 35 adult Long-Evans males were purchased from Charles River Labs and acclimated to the vivarium for at least two weeks before breeding. During breeding, P75–85 females were screened daily for receptive behavior and housed with a breeder male overnight on the day lordosis was observed. All dams were socially housed throughout pregnancy until they were separated into individual cages a few days before giving birth. Day of birth was considered P0.

### Video recording

Home cage behavior recording was conducted with Raspberry Pi 3B+ minicomputers running Debian bullseye with the Raspberry Pi Desktop and equipped with Raspberry Pi Module 1 NoIR cameras. One Raspberry Pi was placed perpendicular to the short end of the cage and closest to the nest location for each cage (**Figure 1**). Cages were set up on wire racks with the water bottle spout facing away from the wall and the camera on the side closest to the wall as rats typically prefer to place their nest near the wall and away from the water bottle. In the event that the dam moved the location of the nest to the opposite end of the cage, the camera side was also switched at the first opportunity. Instances of dams moving the location of the nest to the opposite end during a recording were rare, and those videos were not included. Raspberry Pis were held in place using phone mounts attached to magic arms clamped to the rack and were positioned to capture the width of the front of the cage with a view of the entire cage (except when occluded by excessive bedding or the dam). Pi distance from the cage was not standardized and thus varied slightly between recordings (**Supplemental Figure 2**). Raspberry Pis were programmed to record for one hour starting an hour after lights-off at 30 fps in greyscale at 1280 × 780 or 920 × 550 resolution on postnatal day (P) 0-13. Two infrared LED strip lights (940nm; LED Lights World) were attached to the bottom of the wire shelf above the cages and set to turn on and off for the recording automatically with a digital timer. Another 156 videos in were taken to capture 24-hour recordings at 2 fps and 920 × 550 resolution to capture video footage during both the light and dark phases. These frames were used to improve the pose estimation models, but only videos recorded at 30fps were used to build behavior classifiers. Raspberry Pis were headless and accessed remotely in order to prevent disruption to home cage behavior by experimenter presence before or during the recordings. Following video recording, videos were automatically converted to mp4 format with MP4Box and uploaded to cloud storage. Raspberry Pi recording setup instructions and recording scripts are available at https://github.com/lapphe/raspberry_rat.

### Pose estimation models

Pose estimation for dams was achieved using single animal DeepLabCut^30^. A total of 4,710 frames were extracted from 255 videos. Thirty-two dam body points were selected from 60 candidate body points for labeling based on the user ability to label key points consistently and to provide adequate coverage of the dam regardless of partial body occlusions or orientation relative to the camera (**Figure 1B**). Only body points that were visible in the frame were labeled. For example, if only the left side of the body was visible, points on the right arm, leg, and ventrum were not labeled. Two percent of labeled frames were used for a test set for model evaluation. The dam model was trained for 650,000 iterations using ResNet-50 with a batch size of 8.

The pup pose estimation model was developed using multi-animal DeepLabCut^31^. 1712 frames from 238 videos of pups between postnatal day 0-10 were extracted using k-means clustering and uniform distribution. Nine pup points from the nose to tail base were labeled on each pup visible in the frame (**Figure 1C**). Two percent of labeled frames were used for a test set. The pup model was trained for 200,000 iterations using DLCRNet_ms5 with batch size of 8. Pup detections, rather than individual tracks, obtained by running the DeepLabCut function analyze_videos with auto_track= false, were used for pup pose estimation. A custom python script (PhenoPickleRaw.py) was used to convert the pup detections pickle file (ending in “full.pickle”) to a csv file containing the pup key point detections. Unfiltered dam and pup pose estimation files were joined by frame number using a custom R script (join_dam_pup.py). Column headers were reformatted to match the expected input by SimBA for single animal DeepLabCut pose estimation.

### Behavior classifier development

A SimBA single animal project configuration with a user-defined body points consisting of all dam and pup key points was used for creating behavior classifiers. Because AMBER uses a custom feature extraction script specific to dam and pup points, the only difference between single animal and multi-animal SimBA projects is the expected pose estimation data format imported into SimBA. The width of the wire cage top at the lowest point corresponds to approximately the center of the long side of the cage and was used to define pixels per mm during the Video Settings step (see **Supplemental Figure 2**). Because of the side camera view, the actual pixel/mm distance will change based on the dam and pup location in the cage, but setting this distance helps account for differences in cage distance from the camera and frame resolution between recordings. The outlier correction step of SimBA was skipped.

Dam and pup features were extracted from the pose estimation data using a custom script to calculate 218 features (**Supplemental Table 1)**. Because outlier correction is skipped and a large number of occlusions in each frame is expected for dam key point (e.g. points on the right side are not visible when the left side of her body is facing the camera) and pups (often partially or fully occluded by each other, bedding, or the dam), the majority of feature calculations involve weighting calculations by key point probabilities or applying a minimum probability threshold to exclude occluded points. One feature requires the installation of the circle-fit python package to first fit a circle through the back points then calculate the angle between the first back point, the center of the circle, and the last back point. All other requirements are satisfied by SimBA dependencies.

Classifier training videos were carefully annotated for seven maternal behaviors using BORIS^33^ then imported into SimBA. A total of 3,366,254 frames (31.1 hours of recording at 30 fps) from 28 one-hour videos and 10 additional shorter video clips (1.7-22 minutes each) were annotated for nest attendance, active nursing, passive nursing, licking and grooming, self-directed grooming, eating, and drinking (**Supplemental Table 2**). Clear definitions for inclusion in behavior scoring were developed by incorporating existing definitions for these behaviors, establishing clear rules for the precise start and end of behaviors, and adding and modifying rules until intra-rater and inter-rater reliability was high (>.96) across an initial set of 10 videos for each behavior. To avoid bias in selecting only the most obvious examples of behaviors, the entire recording was scored for all behaviors for each annotated video. The videos were selected for manual scoring included a range of different dams, pups of different ages, and did not include any videos where the nest was at the end of the cage opposite the camera or videos where the dam moved the nest during the recording. The 10 additional video clips were selected to provide more examples of infrequent behaviors (i.e. eating, drinking, passive nursing). A detailed ethogram guide that includes behavior definitions, instructions for scoring, and example images is available at https://github.com/lapphe/AMBER-pipeline.

Random forest models were run in SimBA with the following hyperparameters: 100-1500 trees, 1-2 minimum leaf node, RF_criterion = gini, RF_max_features= sqrt, test size= 20% and no sampling adjustment (**Supplemental Table 2**). Twenty percent of frames were excluded from training and used as a test set to evaluated model performance.

### Pipeline validation for maternal phenotype

Four one-hour recordings were manually scored as described above and used as a hold-out data set to assess model generalizability (not included in the training or test set). These hold-out videos were of four different dams and pups of different ages (P2-9). Passive nursing is an infrequent behavior in Long-Evans rats provided with sufficient bedding material and unfortunately, passive nursing did not occur in any of the four videos precluding evaluation of the passive nursing classifiers in the hold-out video set.

242 videos of one-hour home cage recordings taken beginning one hour after lights off on P1-10 from 49 dams were used in the AMBER pipeline workflow as shown in **Figure 1A** to assess overall patterns of maternal behavior. This set includes the 28 one-hour videos used to create the behavior classifiers. Thresholds used for behavior classifiers are noted in **Supplemental Table 2**. Total duration, percent time, bout number, and mean bout duration were calculated in SimBA for each behavior for each video (**Figure 7** and **Supplemental Figure 5**). Change in behavior durations over time were analyzed using linear mixed models in R with the lmerTest package with litter ID included as a random effect^36^.

### Explanability metrics for behavior classifiers

Feature importance permutations were calculated for each behavior classifier in SimBA. SHAP analysis was run in SimBA on 150 random frames with behavior present and 150 random frames with behavior absent for each model. Full results files for feature importance permutations and SHAP analysis are available on our OSF repository: https://osf.io/e3dyc/.

### Computer hardware and software for machine learning models

All models were trained on a Dell Precision 7920 Tower with a Dual Intel Xeon Gold 5122 3.6GHz processor, 64GB RAM, Windows 10 operating system, and a NVIDIA Quadro P5000 video card. Pose estimation models were trained using DeepLabCut version 2.3. Behavioral classifiers were generated using SimBA version 1.65.5. Python was used for pickle file conversion and pose estimation data joining.

### Data availability

Pose estimation models, scripts, the ethogram guide, and AMBER analysis instructions are available at: https://github.com/lapphe/AMBER-pipeline. Behavior classifier models are available on our OSF repository: https://osf.io/e3dyc/. Please contact the corresponding author with additional inquiries.

## Author contributions

HL collected video recordings, developed the pipeline, analyzed the validation video set, created the figures and tables, and wrote the main text. MS collected video recordings, annotated behavior videos, and edited the text. FC conceived and designed the study, supervised the project, and revised the text. All authors approved the manuscript.

