## Supplemental materials for "Automated Maternal Behavior during Early life in Rodents (AMBER) pipeline"

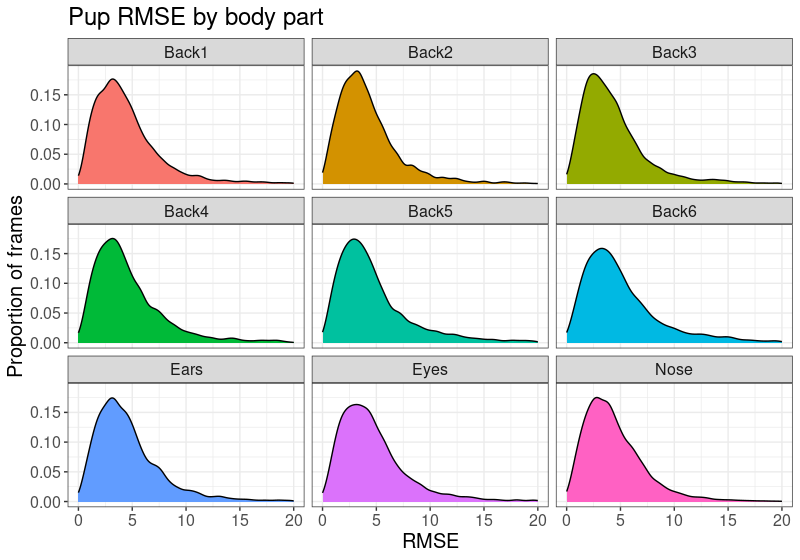

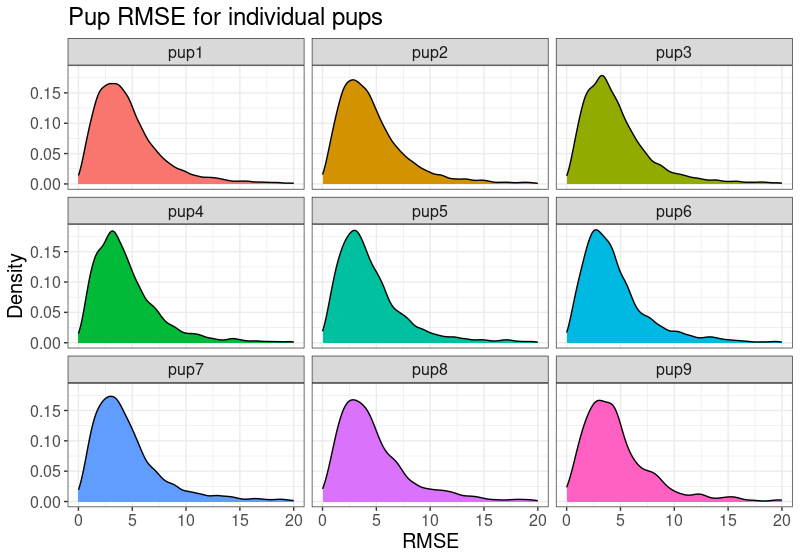

B

A

Supplemental Figure 1. Root mean squared error (RMSE) for pup pose estimation. A) Pup RMSE broken down by pup body point and B) Pup RMSE by individual pup. RMSE was similar across pup body points and individuals.

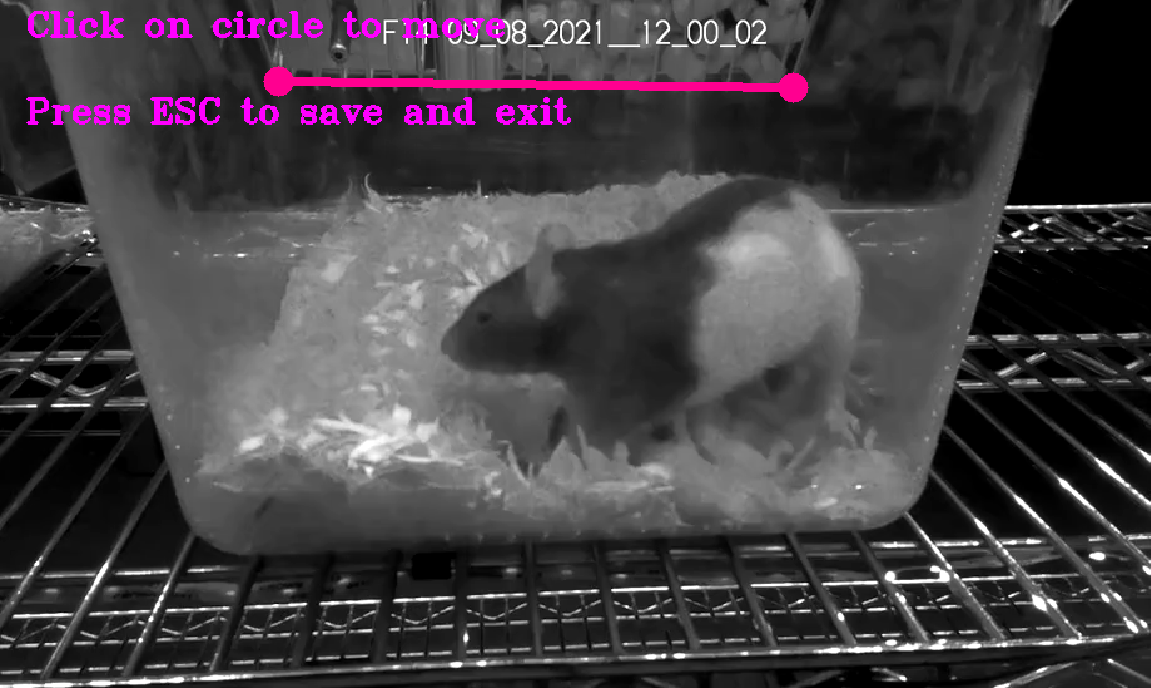

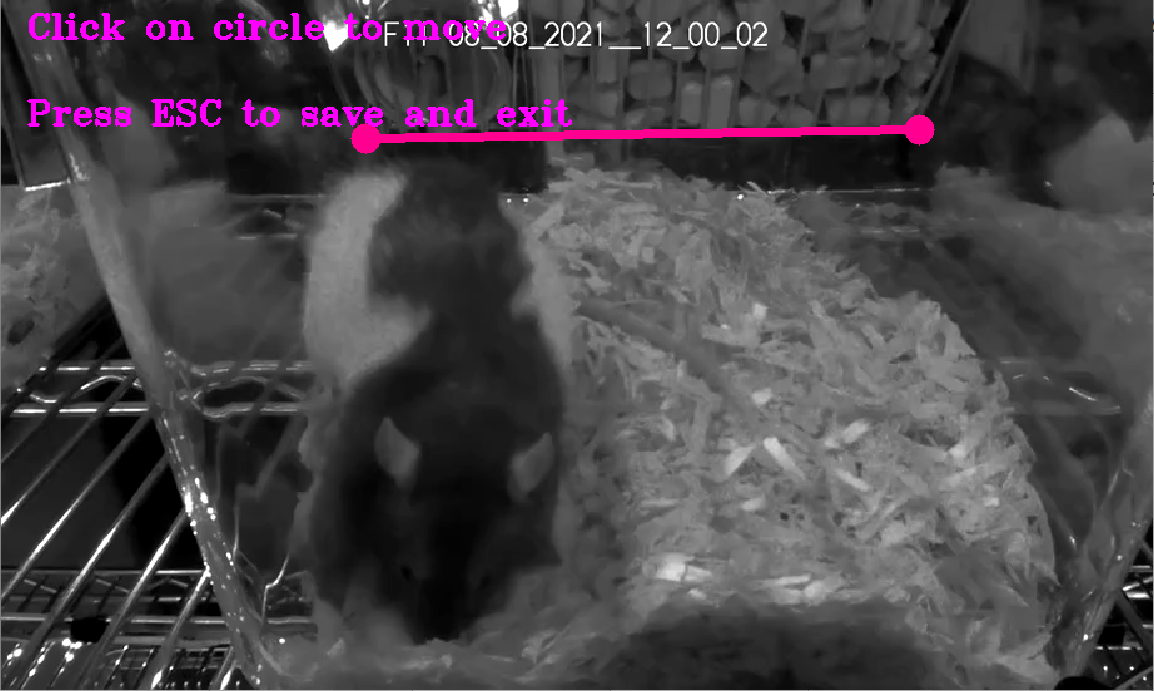

Supplemental Figure 2. Video setting for behavior classifiers. The width of the wire top at the lowest point was used as a reference for determining pixels per millimeter in the videos. This distance is the mid-point of the cage depth. Pixels per mmm is then used during the feature extraction step to normalize some features to account for differences in frame resolution and distance of the camera from the cage during recording.

| **Feature category** | **Description** | **Number of features** | **Features using summary statistics** |
| --- | --- | --- | --- |
| Dam location | Y coordinate of dam head centroid, y centroid, and ventrum | 4 | None |
| Dam-pup distance | Distance between dam body points/centroid and pup centroid | 27 | Sum: 8  Standard deviation: 6  Mean: 8 |
| Dam area | Area of convex hull of all dam point or groups of dam points | 15 | Sum: 3  Standard deviation: 4  Mean: 6 |
| Dam distance | Distances between dam body parts | 12 | Sum: 4  Standard deviation: 2  Mean: 3 |
| Dam angle | Angles between dam body parts | 24 | Sum: 3  Standard deviation: 6  Mean: 10 |
| Pup area | Area of convex hull measures of pup points | 9 | Sum: 3  Standard deviation: 2  Mean: 3 |
| Pup probabilities | Features derived from pup detection probability values | 10 | Sum: 3  Standard deviation:24  Mean: 3 |
| Dam probabilities | Features derived from dam keypoint robability values | 13 | Sum: 4  Standard deviation: 2  Mean: 3 |
| Dam movement | Movement of dam body keypoints or groups of keypoints across frames | 104 | Sum: 30  Standard deviation: 12  Mean: 27 |

**Supplemental table 1.** Summary of custom feature extraction script used for SimBA behavior classification. 217 features were extracted from dam and pup pose estimation data. Features are described based on the data they are derived from (dam, pup, or both), type of measure (coordinate information, areas, distances, and movement), type of calculation (raw number, means, or sums), and the time frame the feature is derived from (within a single frame, between frames [.03s], 0.1s, 1s, or 2s). Numbers refer to the number of features for each variety.

| Behavior classifier | Frames with behavior | Number of trees | Minimum node size | Test set Precision | Test set Recall | Test set F1 | Discrimination threshold | Hold-out set Precision | Hold-out set Recall | Hold-out set F1 |
| --- | --- | --- | --- | --- | --- | --- | --- | --- | --- | --- |
| Nest attendance | 47.8% | 1500 | 1 | 0.996 | 0.997 | 0.997 | .5 | .988 | .993 | .990 |
| Active nursing | 27.5% | 1000 | 1 | 0.982 | 0.970 | 0.976 | .4 | .850 | .808 | .828 |
| Licking & grooming | 11.7% | 1000 | 1 | 0.960 | 0.822 | 0.886 | .38 | .745 | .787 | .766 |
| Passive nursing | 2.5% | 1500 | 1 | 0.966 | 0.976 | 0.986 | .2 | NA | NA | NA |
| Self-directed grooming | 11.2% | 1000 | 2 | 0.981 | 0.826 | 0.897 | .25 | .487 | .634 | .550 |
| Eating | 8.7% | 1000 | 1 | 0.996 | 0.947 | 0.971 | .28 | .517 | .552 | .534 |
| Drinking | 4.4% | 1000 | 1 | 0.991 | 0.991 | 0.927 | .22 | .574 | .535 | .554 |

**Supplemental table 2.** Behavior classifier model training and performance information. Seven random forest behavior classifiers were trained with the model hyperparameters shown. All models achieved excellent performance. The discrimination thresholds shown were applied during analysis of the hold-out video set and maternal phenotype validation set.

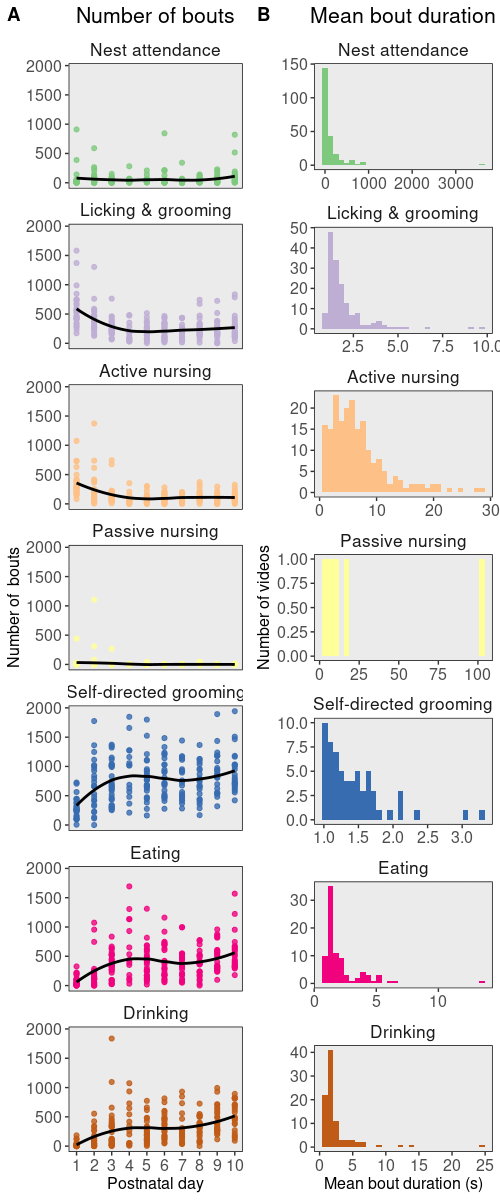

Supplemental Figure 3. Bout number and mean bout duration for maternal phenotype validation video set.
